# Scalable Surrogate Deconvolution for Identification of Partially-Observable Systems and Brain Modeling

**DOI:** 10.1101/2020.03.20.000661

**Authors:** Matthew F. Singh, Anxu Wang, Todd S. Braver, ShiNung Ching

**Affiliations:** Department of Electrical and Systems Engineering, Washington University in St. Louis, St. Louis, MO, USA; Department of Psychological and Brain Sciences, Washington University in St. Louis, St. Louis, MO, USA; Department of Biomedical Engineering, Washington University in St. Louis, St. Louis, MO, USA

## Abstract

For many biophysical systems, direct measurement of all state-variables, *in – vivo* is not-feasible. Thus, a key challenge in biological modeling and signal processing is to reconstruct the activity and structure of interesting biological systems from indirect measurements. These measurements are often generated by approximately linear time-invariant (LTI) dynamical interactions with the hidden system and may therefore be described as a convolution of hidden state-variables with an unknown kernel. In the current work, we present an approach termed surrogate deconvolution, to directly identify such coupled systems (i.e. parameterize models). Surrogate deconvolution reframes certain nonlinear partially-observable identification problems, which are common in neuroscience/biology, as analytical objectives that are compatible with almost any user-chosen optimization procedure. We show that the proposed technique is highly scalable, low in computational complexity, and performs competitively with the current gold-standard in partially-observable system estimation: the joint Kalman Filters (Unscented and Extended). We show the benefits of surrogate deconvolution for model identification when applied to simulations of the Local Field Potential and blood oxygen level dependent (BOLD) signal. Lastly, we demonstrate the empirical stability of Hemodynamic Response Function (HRF) kernel estimates for Mesoscale Individualized NeuroDynamic (MINDy) models of individual human brains. The recovered HRF parameters demonstrate reliable individual variation as well as a stereotyped spatial distribution, on average. These results demonstrate that surrogate deconvolution promises to enhance brain-modeling approaches by simultaneously and rapidly fitting large-scale models of brain networks and the physiological processes which generate neuroscientific measurements (e.g. hemodynamics for BOLD fMRI).

## I. INTRODUCTION

A key challenge in neural engineering pertains to estimating neural model parameters from indirect observations that are temporally convolved from source measurements. For example, many imaging modalities reflect convolution of neural activity with temporal kernels associated with slower physiological processes such as blood flow (Fig. 1A), or molecular concentrations (Tab. I). Often, these kernels are not known, necessitating so-called ‘dual estimation’ of both the latent neural activity and the neural model (including convolutional kernels) at the same time. Our paper presents a computational framework for addressing this problem. Specifically, we assume that the system can be described in the following form (or its discrete-time equivalent):

**TABLE I.** Common neuroimaging measures subject to convolution.

| Modality | Physiology | Popular Kernels | Interpretation |
| --- | --- | --- | --- |
| fMRI | BOLD Signal | Di-Gamma [4] | Hemodynamic Response |
| PET | Radionucleotide Concentration | Multi-Exponential [5],[6] | Exchange, Radio decay |
| $Ca^{2+}$ Imaging | $Ca^{2+}$ Concentration | Multi-Exponential [2],[3] | Diffusion +Kinetics |
| LFP (low freq.) | Membrane Potential | Multi-Exponential/Alpha [7] | $K^+$ Leak +Kinetics |
| Dendritic Recording | Post-Synaptic Potential | Multi-Exponential/Alpha [8],[1] | $K^+$ Leak +Kinetics |

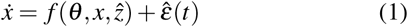

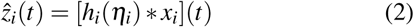

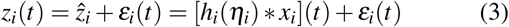

**Fig. 1.**
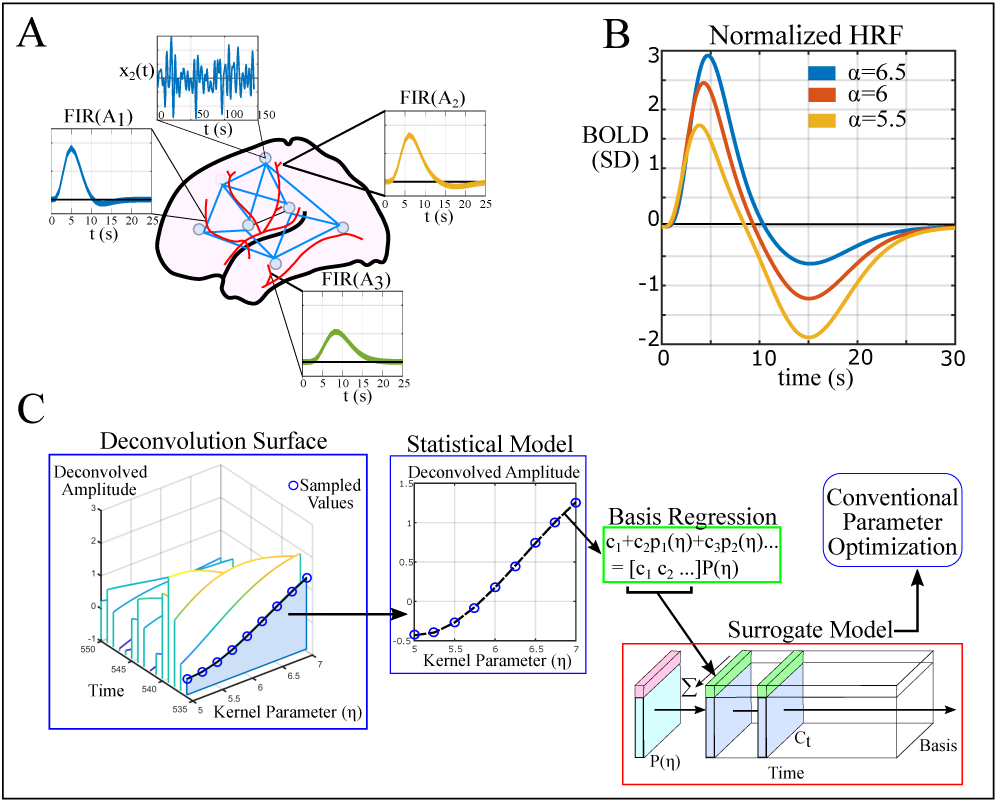
General Approach. A) The brain is an example of a convolutional system when viewed through BOLD fMRI. Dynamics among brain regions are highly nonlinear and usually cannot be directly observed. B) Measurements made using fMRI reflect latent brain activity passing through a hemodynamic response function (HRF). C) Surrogate Deconvolution workflow: a deconvolution surface is estimated by sampling the time-series deconvolution across a variety of kernel parameters (left). A separate surrogate model is formed for each time-point using basis regression to approximate this surface (middle). The combined surrogate models then represent the deconvolution process during parameter estimation.

Here, *x* ∈ ℝ^*n*^ are the hidden non-convolutional state variables and 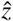 are the physiological variables generated by convolution. We denote unknown parameters for the nonconvolutional plant as *θ* ∈ ℝ^*q*^ and for the convolutional plant as 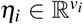. Each parameterized kernel (*h*_*i*_) represents the process generating the corresponding measurable variable *z*_*i*_ via convolution (denoted *). This formulation requires the assumption that these processes may be well-approximated by a finite-impulse response function. We denote process noise in the hidden state variables by 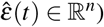 and denote measurement noise *ε*_*i*_(*t*), both of which we assume to be drawn from stationary distributions, independently realized at each time step (noise is not auto-correlated). In the current context, *x* represents latent neural state-variables. The measurements *z*_*i*_ are multi-dimensional recordings of neural data and 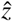 are the corresponding physiological sources. These sources can either feed back into the latent system (e.g. Ca^2+^ concentration) or be modeled as purely downstream (as is typical for BOLD). Formally, we seek to estimate the convolutional kernel parameters {*η*} and the neural model parameters *θ* using the measurements *z* (i.e., the ‘dual’ estimation). This problem formulation is highly relevant to neuroscience and neural engineering since it would enable inferences regarding brain activity via indirect and uncertain dynamical transformations.

### A. Relevance to Neuroscience and Neural Engineering

Whereas many neural models emphasize membrane potentials, channel conductances, and/or firing rate as state variables, high-coverage measurements often consist of the extracellular (“local”) field potential, concentration of signaling molecules (e.g. Ca^2+^), blood-oxygenation (and the derived BOLD-fMRI contrast) or radionuclide concentrations (e.g. PET). In all of these cases, the measurements reflect downstream, temporally extended consequences of neuronal firing (Tab. I). Thus, in the context of neuroscience, the dual-problem consists of simultaneously estimating the parameters of neural systems, while inverting measured signals into their unmeasured neural generators (the state-variables specified by a given model framework). Often this linkage (from generator to measurement) is assumed to be a linear time-invariant (LTI) system so that the relationship can be described via convolution with parameterized kernels. For, example, post-synaptic currents are often modeled via synaptic “kernels” (e.g. “alpha-synapse”, [1]), kernels for molecular concentrations (e.g. Ca^2+^, [2],[3]) are derived from Markovian kinetic-schemes ([9],[2]), and the neurovascular coupling kernel (linking BOLD-fMRI and neural “activity”) is described by a Hemodynamic-Response Function (HRF, [4]; Fig. 1A,B,). If these functions are assumed fixed, it may be possible to estimate the neural state-variables via deconvolution, in which case, conventional modeling approaches are feasible. However, in many cases, only the general form of the kernel function is known (e.g. up to a small number of unknown parameters) resulting in computationally difficult dual estimation problems (estimating the neural states and the model parameters). The current work aims to treat such dual problems in a computationally efficient, and highly scalable manner.

### B. Previous Work

Currently, there are several methods to deal with dual-identification for small systems and these approaches may be grouped into black-box and grey box models. However, whereas black-box modeling encompasses diverse approaches such as neural networks ([10]), Volterra Expansion ([11]), and Nonlinear Autoregressive Moving Average (NARMA) models ([12]); grey-box identification (model parameterization) has largely centered upon the dual/joint Kalman-Filters (linear, extended, unscented etc. [13], [14], [15], [16]) and related Bayesian methods. Under these approaches, the convolutional component is converted into the equivalent linear time-invariant (LTI) system format and free-parameters are modeled as additional state-variables. Thus, joint state-space techniques re-frame the dual-estimation problem as conventional state-estimation with a fully-determined model.

However, none of these methods are well situated to perform dual-identification for large (grey-box) systems due to the high computational complexity and data-intensive nature of Bayesian dual-estimation. These features are particularly limiting to neuroscience applications which typically feature a large number of connectivity parameters and potentially few sampling times (e.g. fMRI). These approaches also increase in complexity with the number of additional state variables necessary to represent complex kernels such as the hemodynamic response function (Fig. 1B).

Neural systems present two challenges to the current status quo: the dimensionality of the neural system/parameters and the complexity of the convolutional kernel. Neurobiological recordings are often high-dimensional, containing dozens or hundreds of neurons/neural populations. Moreover, the number of free parameters often scales nonlinearly with the number of populations (e.g. quadratically for the number of connectivity parameters). Current dual-estimation approaches such as joint-Kalman-Filtering are computationally limited in these settings due to their high computational complexity in terms of both the number of state variables and the number of parameters estimated.

Previous approaches are also limited in terms of kernel complexity. Since joint-Kalman-Filtering employs a state-space representation, convolutional variables are implicitly generated via linear time-invariant (LTI) systems. This issue is not inherently problematic, as many neural models contain simple exponential kernels which are easily converted to an additional LTI variable (e.g. Tab. I). However, specific domains feature higher-order kernels such as the Hemodynamic Response Function (HRF) that relates latent neural activity and observed BOLD signal in fMRI. Approximating the HRF through a linear time-invariant (LTI) system requires multiple additional layers of state-variables which greatly increases the difficulty of estimating neural activity and also increases the overall computational burden.

## II. Approach

We propose to treat this problem by directly performing optimization within the latent state-space using Surrogate Models to replace the state-estimation step (Fig. 1C). Surrogate functions comprise a means to approximate computationally intensive functions, typically through a linear combination of *a priori* specified nonlinear bases (e.g. polynomial families, radial-basis functions etc.). In the current case, we propose using surrogate models to explicitly estimate latent variables by deconvolving the measured time-series by the current estimate of the convolutional kernel at each iteration. Deconvolution is typically performed either by iterative algorithms such as the Richardson-Lucy algorithm ([17],[18]), Alternating Direction Method of Multipliers (e.g. [19]; ADMM) or explicit transformations in the Fourier domain. The proposed surrogate techniques are compatible with any combination of deconvolution algorithm and additional signal processing that are smooth with respect to the kernel parameters. In a later example with empirical fMRI data, we use the Wiener-deconvolution ([20]) coupled with variance normalization in the time-domain:

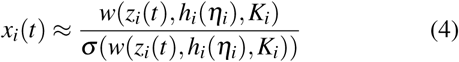

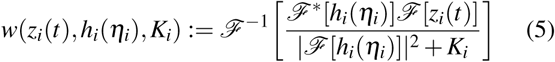

with *w*(*z*_*i*_, *h*_*i*_(*η*_*i*_), *K*_*i*_) denoting the Wiener deconvolution of *z*_*i*_ with respect to kernel *h*_*i*_(*η*_*i*_) and noise-factor *K*_*i*_ equal the mean power-spectral density of the measurement noise *ε*_*i*_(*t*) divided by the mean power spectral density of *z*_*i*_. We denote standard deviation by *σ* and *ℱ, ℱ*^*^ denote the Fourier transform and its complex conjugate, respectively. Through deconvolution, we reduce the dual-estimation problem to conventional system identification with the convolutional kernel as an additional free parameter. Using surrogate models we reduce deconvolution-algorithms into simple, differentiable functions of the kernel parameters (Fig. 2A). Thus rather than solving the dual estimation problem:

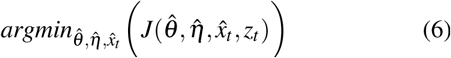

for some loss function *J*, we solve the parameter-estimation problem:

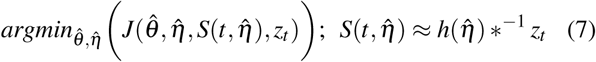

for which *S* denotes the Surrogate Deconvolution model and *^−1^ is a user-defined deconvolution algorithm, potentially incorporating priors on the distribution of *ε*_*i*_(*t*) and further signal processing (e.g. normalization or additional filtering). In later experiments, we set *J* as the mean-squared error of 1-step predictions, e.g.

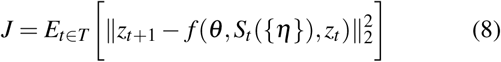

with *T* denoting the set of initialization times during training. The surrogate model *S* is a linear combination of smooth, nonlinear bases and is therefore smooth for both iterative deconvolution algorithms, such as Richardson-Lucy, and for explicit transformations. Thus, algorithms which are natively nonsmooth due to randomization or stopping criteria (e.g. Richardson-Lucy) are converted to an accurate, but differentiable form via the Surrogate representation (e.g. Fig. 2B). The remaining, (surrogate-assisted) fitting problem is thus amenable to highly scalable techniques such as gradient-based optimization.

**Fig. 2.**
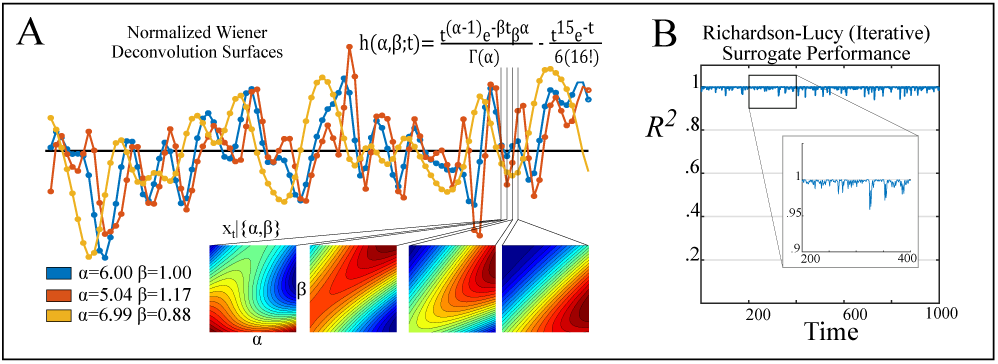
Surrogate surfaces for performing deconvolution. A) Example fMRI time-series deconvolved by hemodynamic models with different values for the kernel parameters (*α, β*). The inset equation is the hemodynamic response function (kernel). Subplots show the surrogate surfaces for four sequential time-points (e.g. the deconvolved signal amplitude at that time as a function of kernel parameters). Note that the surrogate surfaces at a fixed time-point are smooth, whereas the variation in deconvolved signal across time is much less regular. B) Performance in reconstructing the iterative Richardson-Lucy deconvolution of the same signal across an 81×81 grid (same range as A) using third-order bivariate polynomials (Appendix Sec. VI-C). Inset shows a representative stretch of 200 points (144s).

### A. Contributions

Our contribution in this regard is generating surrogate models to explicitly approximate the deconvolution process in a computationally-efficient closed form. To our knowledge, previous approaches have not sought to estimate nonlinear models using parameterized deconvolution. We do so in a two-step process. First, we build a surrogate model of the deconvolution process (deconvolving *z*_*i*_(*t*) by *h*_*i*_(*η*_*i*_) as a direct function of the kernel parameters *η*_*i*_). We fit one surrogate function per measurement in the deconvolved space: the value of a deconvolved channel evaluated at a specific time. For a fixed basis, this representation may be fit rapidly at scale. For example, the empirical brain data treated later requires nearly two million surrogate functions per subject (419 brain regions × 4444 time points), all of which can be fit in seconds as the only computation of nonlinear complexity is shared across time points (Eq. 9). In the second step, we directly integrate the surrogate model into optimization algorithms. By doing so, the time-course of each latent state-variable is expressed as a direct, easily differentiable, function of the kernel parameters (*η*).

We present these results as follows. First we introduce surrogate methods and the proposed technique, Surrogate Deconvolution, in which surrogate models of the latent variable are directly integrated into the optimization procedure. In the next section we consider the special case of gradient-based optimization and demonstrate how error-gradients are efficiently back-propagated through the Surrogate Model. We then test Surrogate Deconvolution in two sets of experiments. First, we consider a low-dimensional case (a small LFP model) in which existing techniques for dual-estimation remain tractable. This simplified setting allows us to benchmark Surrogate Deconvolution’s accuracy in parameter/state estimation relative the joint-Extended Kalman Filter and the joint-Unscented Kalman Filter. Results demonstrate that Surrogate Deconvolution is competitive even within the Kalman Filter’s operating domain. We then consider more complicated fMRI models in which current dual-estimation techniques are not applicable due to high-dimensionality and kernel complexity. We demonstrate that Surrogate Deconvolution is robust to spatial variation in the HRF kernel in contrast to state-of-the-art non-dual approaches. Lastly, we demonstrate the approach’s feasibility to empirical fMRI data. Thus, Surrogate Deconvolution performs competitively within the scope of current dual-estimation approaches and enables robust dual-estimation for a much larger set of problems than previously considered.

## III. Surrogate Deconvolution

Our procedure contains two parts. First, we construct a surrogate function for each channel and time-point, a process which can be massively parallelized, if necessary. We use the surrogate construction to express the estimation of unobserved state variables as a direct function of *η*. The surrogate function then replaces unobserved variables in a user-chosen identification-algorithm for fully observable systems. This process is advantageous as it enables direct calculation of how changing parameters *η* influence the final estimate of unobserved state variables (for the current set of parameters) as opposed to techniques such as the dual Kalman Filter which do not “look-ahead” to see how changing downstream parameters will affect state estimates (since ▽_*η*_*f* = 0 without a surrogate model).

The key insight underlying surrogate deconvolution regards the effect of varying a kernel parameter. As demonstrated in Fig. 2, changing a kernel parameter produces intricate effects upon deconvolved estimates when viewed from the time-domain. Even when these effects can be expressed analytically (as in the Wiener deconvolution) they are not readily reduced to a temporally-local calculation using first-principles. However, when the kernel is lower frequency than the signal (as usually happens in biology), the effect of kernel variation on a single estimate is often quite smooth with respect to the kernel parameter. Thus, the effect of kernel variation on a single deconvolved estimate is very well-approximated by simple functions of the kernel parameter. Together, these functions comprise the surrogate model.

### A. Building Surrogate Representations

We efficiently define and evaluate surrogate models by storing coefficients in tensor format. For a vector of *m*_*i*_ stacked basis functions 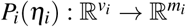 we define the 3-tensor *C* defined for each channel (“i”) and a prior distribution on *η*:

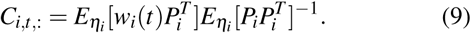

Thus, *C* stores the coefficients of regressing the basis functions *P*_*i*_ on the deconvolved time series *w*_*i*_ (one of *P*_*i*_’s bases should be [*P*_*i*_] _*j*_ = 1, ∀*η*_*i*_ to provide the intercept). For clarity of presentation, we have reduced the input arguments of *w*_*i*_ to time alone. By *E*_*ηi*_ we denote the expectation taken over some prior distribution on *η*_*i*_. In practice, the choice of prior is not usually impactful, as an arbitrarily fine sampling of the response surface can be quickly computed in parallel and the surrogate goodness-of-fit can be similarly increased by adding additional (linearly independent) basis functions. In all later examples, we simply assume a uniform distribution over reasonable bounds. The tensor *C* holds coefficients of each time-point’s surrogate model with *C*_*i,t, j*_ denoting the coefficient of basis *j* in predicting the deconvolution of channel *i* at time *t* in the deconvolved time-domain (which is shifted from the measurement times). We evaluate the surrogate functions in parallel by defining the following product between 3-tensor *C* and a 2-tensor-valued function [*P*({*η*})]_*i, j*_ := [*P*_*i*_(*η*_*i*_)] _*j*_:

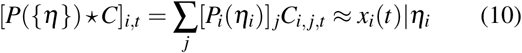

with the right-hand side denoting the optimal estimate of *x*_*i*_(*t*) (e.g. in the least-squares sense for Wiener deconvolution) given *η*_*i*_, *z*_*i*_(*t*) and any fixed priors used to define the chosen deconvolution. In principle, this technique could be used for system identification objectives in which errors are defined in terms of predicting *x*_*t*_ or *z*_*t*_ or both. In practice, however, we have found that including *x*_*t*_ predictions within the objective function leads to a moving-target problem in which identification algorithms enter periods of attempting to maximize auto-covariance (by changing *η*). Therefore, we assume that objectives are given of the form:

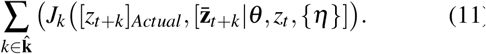

Here, 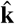 denotes the user-determined time steps at which to evaluate the cost function *J* which potentially varies by time-step (e.g. choosing to weight temporally distant predictions less). The right-hand side denotes the current estimate 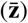 of *z*_*t*+*k*_ given *θ*,{*η*}, and *z*_*t*_. Thus, the new cost function incorporates the actual measurements and their prediction. However, unlike conventional dual approaches, the predictions are a direct, explicit function of previous measurements, rather than in terms of both measurements and an iteratively estimated latent variable.

### B. Deploying Surrogate Models

To evaluate the cost function, we use make forward predictions in the latent-variable (deconvolved) domain and then convolve those predictions forward in time to evaluate error in terms of observations. For *k*-step predictions and kernel length *τ*, this corresponds to:

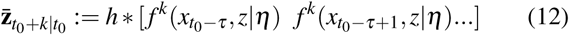

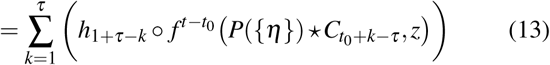

We use 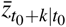 to denote the estimate of *z*_*t*+*k*_ using initial conditions for the convolutional variable (*z*) and latent variable (*x*) prior to *t*_0_. The operator ∘ denotes the Hadamard product (element-wise multiplication). In the latter equation, we have condensed notation for the effect of *z* on *f* as follows:

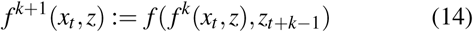

with *f*(*x*_*t*_, *z*) := *f*(*x*_*t*_, *z*_*t*_). Thus, *f*^*k*^ is not a proper iterated composition when it accepts both *x*_*t*_ and *z*_*t*_ as arguments, since only one variable (*x*_*t*+1_) is output. However, we abuse this notation for clarity of presentation. For brevity, we also use 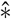 to indicate convolution over initial conditions as indicated in the variable indices. Hence the earlier equation (Eq. 13) condenses to:

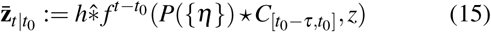

As a general technique for re-representing dual estimation problems, Surrogate Deconvolution is compatible with most estimation techniques. However, the approach is particularly advantageous for gradient-based estimation as the deconvolution process is replaced with an easily differentiable surrogate form. For single-step prediction, the resulting error gradients for the nonlinear plant’s parameters (*θ*) and the convolution kernel parameters ({*η*}) are as follows:

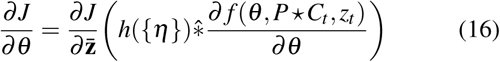

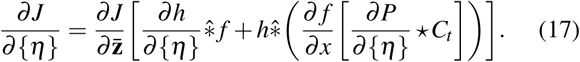

Thus, surrogate deconvolution re-frames dual-estimation problems into conventional parameter-estimation problems which are amenable to gradient-based approaches. The analogous gradients for multi-step prediction are derived by augmenting the one-step prediction gradients with back-propagation through time. We demonstrate the power of surrogate deconvolution by reconstructing large brain network models from either simulated data or empirical recordings.

## IV. Data-driven Model Identification

We present two applications to brain discovery to illustrate the advantage of surrogate deconvolution-enhanced methods for both state-estimation (Kalman-Filtering) and grey-box parameter identification. Both examples are dual-estimation problems (state and parameter), but we assess their performance in the state and parameter components separately to make comparisons with existing work which may be particularly designed for either domain. For instance, dual-estimation using the joint unscented Kalman Filter has been particularly successful in parameterizing black-box models for filtering (e.g. [15]), but requires further modification in some more complicated grey-box models for which the conventional Kalman gain (1-step prediction) is provably rank-deficient. To demonstrate the adaptability of surrogate deconvolution we consider two different simulated system identification /estimation problems and one empirical application.

### A. Modeling and Isolating Local Activity from the LFP

Our first example compares performance across methodologies designed for state-estimation. This simulated problem consists of identifying the wiring of a neural system and subsequently reconstructing cellular activity from simulated extracellular recording of the “local” field potential (LFP). This signal is primarily generated by the combined currents entering into the local population of nerve cells as opposed to the currents directly generated by neural firing or the trans-membrane potential. Thus, the measured LFP reflects the temporally extended effects of input into a brain region rather than the current population activity (Fig. 3A). To describe this process, we use a three-level discretized-model combining 10 coupled neural-mass models (*n*_*pop*_ = 10) with passive integration of post-synaptic currents:

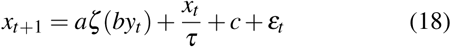

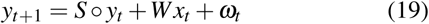

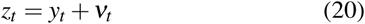

**Fig. 3.**
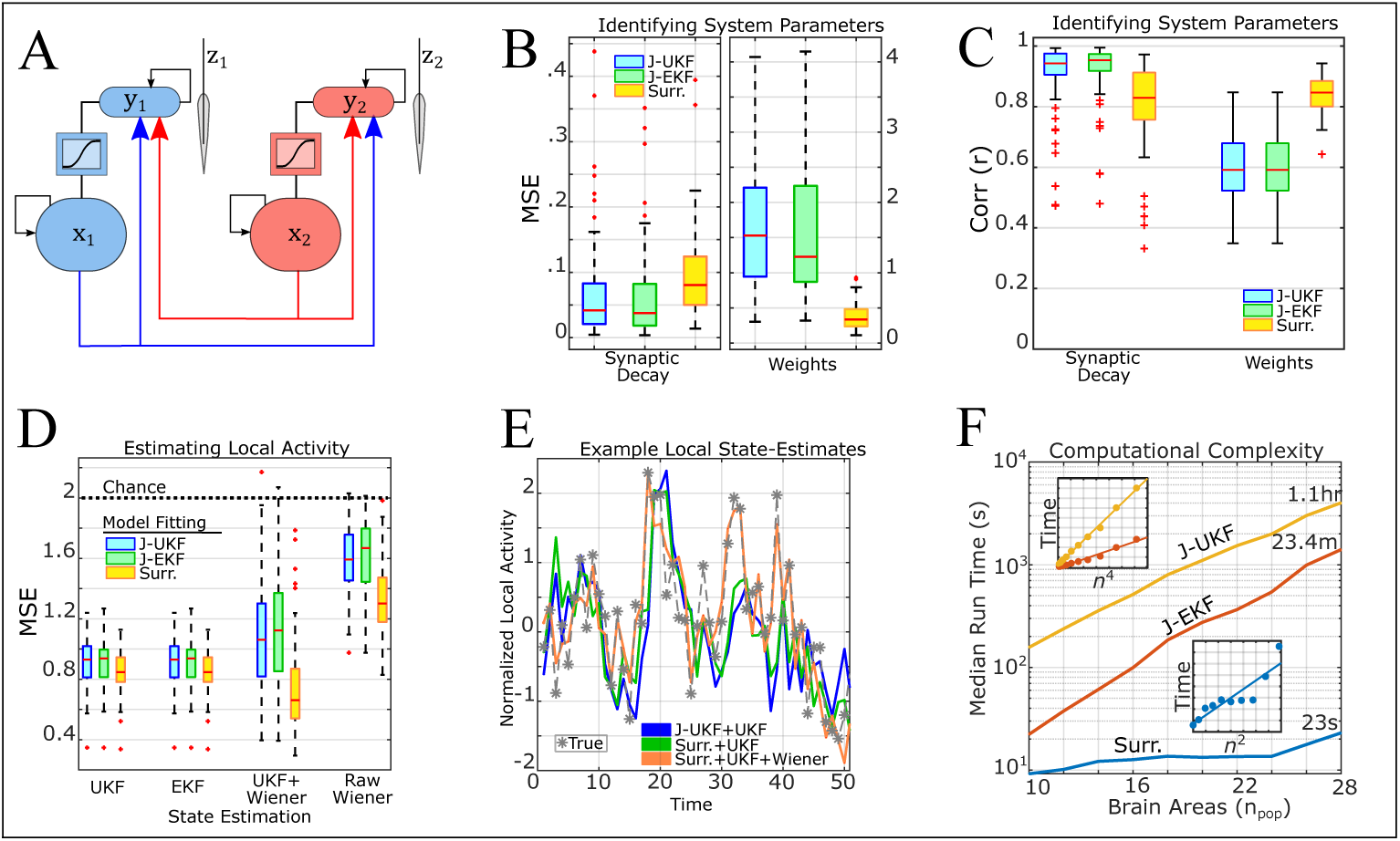
Surrogate deconvolution’s performance in inverting a neural-mass model of Local Field Potential. A) Model schematic: output signals from each population arrive at post-synaptic terminals. Electrode measurements primarily reflect the post-synaptic potentials generated from synaptic activity. B) Bench-marking total error in identifying the synaptic time-constants and network connectivity. Surrogate deconvolution is compared to the current gold-standard: joint-Kalman Filters (Unscented and Extended). C) Same as (B), but displaying performance in terms of correlation rather than mean-square-error. D) Performance in reconstructing local spiking activity from electrode measurements using the identified system models for a variety of state-estimation techniques. E) State estimation performance for a representative case (the simulation with median jUKF+UKF performance). F) Computational complexity of system identification methods in terms of the number of brain regions considered. The top inset shows the run-time and least-squares fit on rescaled x-axes 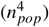 to demonstrate the 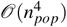 complexity of Kalman-Filtering in terms of the number of brain regions (*n*_*pop*_). The bottom inset is the same, but for surrogate deconvolution and 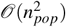. Calculations were performed single-core on an Intel Xeon E5-2630v3 CPU.

Here, *x* is the synaptic-gating variable which describes neural activity. The sigmoidal transfer-function is denoted *ζ* (*x*) := (1 + *exp*[−*x/*5])^−1^ with scaling coefficient *a* = 3. The time constant of *x* is denoted *τ* and baseline drive to *x* is denoted *c*. The parameters *a, τ* ∈ R and 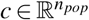 are assumed known as are the covariances of process noise *ε*_*t*_, *ω*_*t*_ and measurement noise *v*_*t*_ (see Appendix). Thus, the unknown parameters are the connections between neural populations (*W*) and the synaptic time-constants *S*. We considered two general approaches to system identification: either using the current gold-standard (joint Kalman estimation) or using surrogate deconvolution for least-squares optimization. The joint Kalman filter and associated variants operate identically to the original Kalman filter, except that the state-space model is augmented with unknown parameters being treated as additional state-variables with trivial dynamics (e.g. 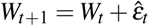 and similarly for *S*). The “noise” terms 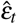 for parameter state-variables are assumed iid. and with a user-defined variance that determines the learning rate. Based upon numerical exploration, we found that the best performance for both EKF and UKF was with an initial prior on parameter variance 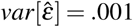. Every 50 time-steps we decreased the variance prior by 5% of its current value.

For comparison with existing techniques we used both the joint-Extended Kalman Filter (jEKF) which linearizes the nonlinear portion of dynamics and the joint-Unscented Kalman Filter (jUKF) which directly propagates noise distributions through nonlinearities using the Unscented Transformation ([14]). We compare these traditional methods with system identification through surrogate deconvolution. The benefit of surrogate deconvolution is the ability to apply a wide variety of optimization techniques to partially-observable identification problems which can decrease computation time over conventional techniques (Fig. 3F) and expand the scope of problems which may be tackled. For this first example, we have chosen a relatively simple case (low-dimensional, single-exponential kernels) so that conventional methods (Kalman Filtering). Therefore, the goal of this test is not to demonstrate an overwhelming advantage of surrogate deconvolution over Kalman Filtering, but to show that the proposed technique can perform competitively in cases for which Kalman-Filtering is applicable, but imperfect. Subsequent examples will consider cases in which Kalman Filtering is not tenable.

To implement Surrogate Deconvolution, we first reformulate this problem as a convolutional equation through the change of variable *r*_*t*_ := *Wx*_*t*_ :

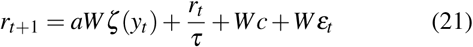

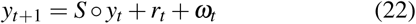

or, equivalently,

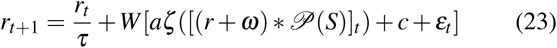

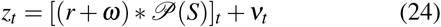

with *𝒫*(*S*) denoting the discrete-time kernel formed from polynomials of *S* to a suitably long length [0 1 *S S*^2^ *S*^3^*…*] analogous to exponential decay for continuous-time systems. In this form, the parameters can be estimated using traditional least-squares methods, optimizing over *W* and *S*. However, by leveraging the tensor representation of surrogate models, this equation can be reduced into a single equation in *S* by representing the optimal choice of *W* for a given *S* as a direct function of *S*. To do so we define the matrix

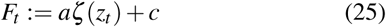

and the associated 3-tensor

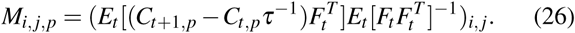

Each *n* × *n* page of this tensor (e.g. the matrix formed by holding *p* constant) stores the coefficients of the least squares solution for *W* in predicting *C*_*t*+1,*p*_ − *C*_*t,p*_*τ*^−1^ from *F*_*t*_ for the *p*^*th*^ basis function. Since 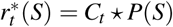, for a given synaptic decay term *S*:

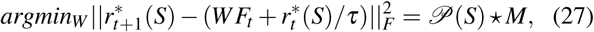

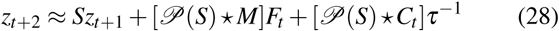

with *r*^*^(*S*) denoting the estimate of *r* produced through the surrogate deconvolution of measurements *z* with the kernel *𝒫*(*S*). Thus, in this case, surrogate deconvolution enables the approximation of 2*n*_*pop*_ difference equations containing *n*_*pop*_(*n*_*pop*_ + 1) unknown parameters (*W* and *S*) to *n*_*pop*_ difference equations with *n*_*pop*_ unknown parameters (only *S*).

The resultant model (from Eq. 28) is also compatible with a wide variety of optimization techniques. For simplicity, we fit the parameters *S* through ordinary least-squares optimization in terms of predicting *z*_*t*+2_ as in Eq. (28). Optimization was performed using Nesterov-Accelerated Adaptive Moment Estimation (NADAM; [26]) with both NADAM memory parameters set equal to .95, and the NADAM regularization parameter set to .001. Training consisted of 15,000 iterations with each minibatch containing 1000 time points. The step size (learning rate) of updates was .0001.

All methods were able to retrieve accurate estimates of the synaptic decay term *S* (Fig. 3B,C). The best-performing method varied by simulation (e.g. for different true values of *W, S*), but the mean error was greater for surrogate deconvolution than Kalman Filtering methods (Unscented and Extended) which performed near-identically. By contrast, surrogate deconvolution always provided a more accurate estimate of the connectivity weight parameter (*W*) and the advantage relative Kalman-Filtering was substantial (Fig. 3B,C). The poor performance of the Kalman Filter for identification in this case is not surprising as the Kalman Filter is known to be non-robust for large systems ([24]) and the *W* parameter adds 100 additional latent state-variables to the joint Kalman model as opposed to the 10 state-variables added by *S*.

### B. Reconstructing Firing-Rate from LFP

We next examined the ability of each method to recover the time series of neural activity *x*_*t*_ using the previously generated state-space models. During this stage, models produced during the previous identification step were used to estimate the latent state variable *x*_*t*_ (Fig. 3D,E). It is important to distinguish between state-estimation techniques (e.g. UKF) which we used to estimate *x*_*t*_ from previously-fit models and the techniques used to fit those initial models (e.g. jUKF) as these steps need not “match” (e.g. UKF-based state-estimation from a surrogate-identified model). Measurements consisted of simulated extracellular voltages *z*_*t*_ generated by resimulating the same ground-truth model (i.e. the same parameters, but new initial conditions and noise realizations). As in the identification stage, we considered two general approaches to recovering the latent variable *x*_*t*_ : either through deconvolution or Kalman Filtering (unscented and extended). Kalman filtering in this setting produces direct estimates of *x*_*t*_ and *y*_*t*_. By contrast, deconvolving *z*_*t*_ ≈ *y*_*t*_ produces an estimate of *r*_*t*_, so deconvolution estimates of *x*_*t*_ were produced by premultiplying the deconvolved time series with 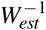 (the inverse estimated connectivity parameter). We considered deconvolution applied either directly to the raw measurements (*z*_*t*_) or to the estimates of *y*_*t*_ produced by Kalman filtering *z*_*t*_ with the estimated models (both unscented and extended Kalman filters were considered). Noise covariance estimates for Kalman filtering at this stage were the same as those assumed in the initial stage: a value close to the mean tendency, rather than the true values which were randomly selected for each simulation.

We found that the type of Kalman Filter used for state-estimation had no appreciable effect upon accuracy (Fig. 3D). Likewise, the technique used for system identification (surrogate deconvolution vs. EKF/UKF) had little effect, although surrogate deconvolution was slightly more accurate on average. However, model performance differed greatly for deconvolution-based state-estimation (using *x*_*t*_ ≈ *W*^−1^[*𝒫*(*S*) *^−1^ *y*_*est*_]_*t*_). Models estimated using joint-Kalman Filtering (jEKF/jUKF) performed worse using deconvolution-based estimation of *x*_*t*_ than Kalman-based estimation (Fig. 3D). This result is unsurprising as the deconvolution-based estimate additionally requires the inverse weight parameter *W*^−1^ and both jUKF and jEKF poorly estimated *W*. Interestingly, however, estimation accuracy for surrogate-identified models decreased when using deconvolution of the raw, unfiltered measurements, but increased for the UKF+decconvolution hybrid. The former result is not surprising as pure deconvolution is clearly suboptimal in not considering the noise covariance. This result was unexpected and it suggests the possible benefit of using a two-stage estimation procedure in which Kalman-Filtering first dampens measurement noise and improves estimates of measurable state-variables. Then, subsequent deconvolution might improve the estimate of latent state-variables by considering the impact of estimates across time, rather than just the directly subsequent measurement. However, these benefits are likely situation-dependent and therefore require more study. In any case, results indicate that state-estimates from models produced by surrogate-deconvolution are at least as accurate as those produced by Kalman-Filtering and potentially more so depending upon the state-estimation procedure (Fig. 3D,E).

### C. Computational Efficiency

Surrogate deconvolution is also computationally efficient as it scales linearly with the number of measurement channels (*O*(*n*)) in both forming and evaluating surrogate functions which is also parallelizable across channels. However, since Surrogate deconvolution is not a system identification procedure in and of itself, time-savings depend upon how the technique is used (e.g. which optimization scheme it is coupled to). The advantage of surrogate deconvolution is that it can be combined with a wide-variety of optimization techniques which are otherwise ill-suited to partially-observable problems. In this first simulation, for instance, the number of parameters scale with *n*_*pop*_(*n*_*pop*_ + 1) and the number of state variables (in the native space) scale with 2*n*_*pop*_. Thus, the dominant complexity of joint-UKF and joint-EKF is greater than 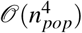 as joint-UKF/EKF are *𝒪*(*n*^2^) in the number of parameters and at least *𝒪*(*n*^2^) in the number of native (non-parameter) state-variables. By contrast, the gradient approaches applied with surrogate deconvolution have approximately 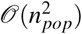 complexity (Fig. 3F). However, surrogate deconvolution is not limited to gradient-based approaches. The main effect is to simplify error functions to a direct equation in the measurable variables so surrogate deconvolution is compatible with a wide variety of non-gradient techniques, as well (e.g. heuristic-based or Bayesian). As such, surrogate deconvolution presents the opportunity to identify significantly larger partially-observable systems than previously considered.

### D. Reconstructing Connectivity and Hemodynamics in Simulated fMRI

For our second example, we considered the ability to correctly parameterize large-scale brain models from simulated fMRI data. Brain regions were modeled through the continuous-valued asymmetric Hopfield model ([25]) and simulated fMRI signals were produced by convolving the simulated brain activity with randomly parameterized Hemodynamic Response Function (HRF) kernels ([4]):

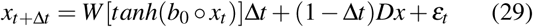

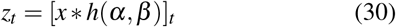

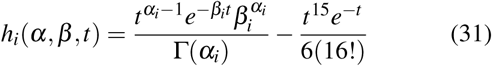

Parameter distributions for simulation are detailed in the Appendix. Simulations were integrated at Δ*t*=100ms and sampled every 700ms (mirroring the Human Connectome Project’s scanning TR of 720ms [27]). Simulated HRF’s (*h*_*i*_) were independently parameterized for each brain region according to the distributions *α*_*i*_ ∼ *𝒩* (6, *σ* ^2^) and *β*_*i*_ ∼ *𝒩* (1, (*σ/*6)^2^) in which the variability term *σ* was systematically varied. Each HRF can be well approximated by a finite-length kernel and therefore can be represented as a discrete-time linear plant. However, doing so, in this case, requires multiple hidden state-variables per region which impairs Kalman-based dual-estimation procedures. Instead, most current procedures to deal with fMRI-based systems identification at scale ignore inter-regional variability in *h*_*i*_ and instead seek to retrieve *x*_*t*_ by fixing HRF parameters (e.g. [31], [30]) to the so-called “canonical HRF” (e.g. *α* = 6, *β* = 1). In this example, we demonstrate the potential pitfalls of this assumption and the benefits accrued by efficiently fitting hemodynamics through Surrogate Deconvolution. To do so, we attempted to reconstruct *W* using Mesoscale Individualized NeuroDynamics (MINDy) in either its base form (which assumes a canonical HRF) or in an augmented form in which the predictions are calculated as in Eq. 13. MINDy model fitting consists of using NADAM-enhanced gradient updates ([26]) to minimize the following cost function:

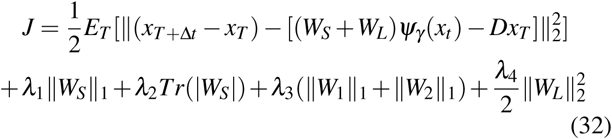

in which the estimated weight matrix *Ŵ* is decomposed into the sum of estimated sparse (*W*_*S*_) and low-rank (*W*_*L*_) components satisfying:

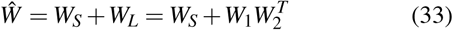

for some *W*_*S*_ ∈ 𝕄^*n*×*n*^ and *W*_1_,*W*_2_ ∈ 𝕄^*n*×*k*^. The hyperparameter *k* < *n* determines the rank of the low-rank component *W*_*L*_ and the regularization hyperparameters {*λ*_*i*_} define statistical priors on each of the weight matrix components (Laplace prior for *W*_*S*_, *W*_1_,*W*_2_ and a normal prior for 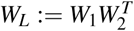). This decomposition has been shown useful to estimating large brain networks ([30]). The nonlinear function *ψ* is parameterized by the parameter vector *γ* ∈ R^*n*^ with

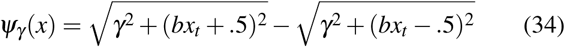

For the Surrogate-Deconvolution, however, these analyses are performed in the original space to prevent the aforementioned moving target problem. Hyper-parameter determination and simulation parameters are detailed in the appendix.

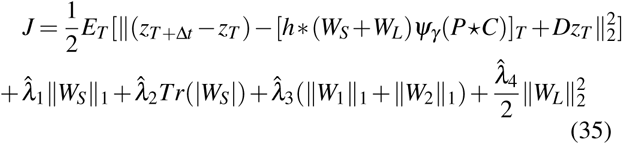

Results demonstrated a clear benefit for additional modeling of the local hemodynamic response (Fig. 4A,B). When hemodynamics differed only slightly between simulated brain regions both methods produced highly accurate estimates of the connectivity parameter *W*. However, past *σ* = .4 (the SD of spatial variation in one of the HRF parameters), the accuracy of estimated connectivity rapidly decreased for conventional methods, while only slightly decreasing for surrogate deconvolution. In addition, the hemodynamic parameter estimates also became increasingly accurate. Thus, surrogate deconvolution enables accurate system (brain) identification when the downstream plants (hemodynamics) are variable and unknown.

**Fig. 4.**
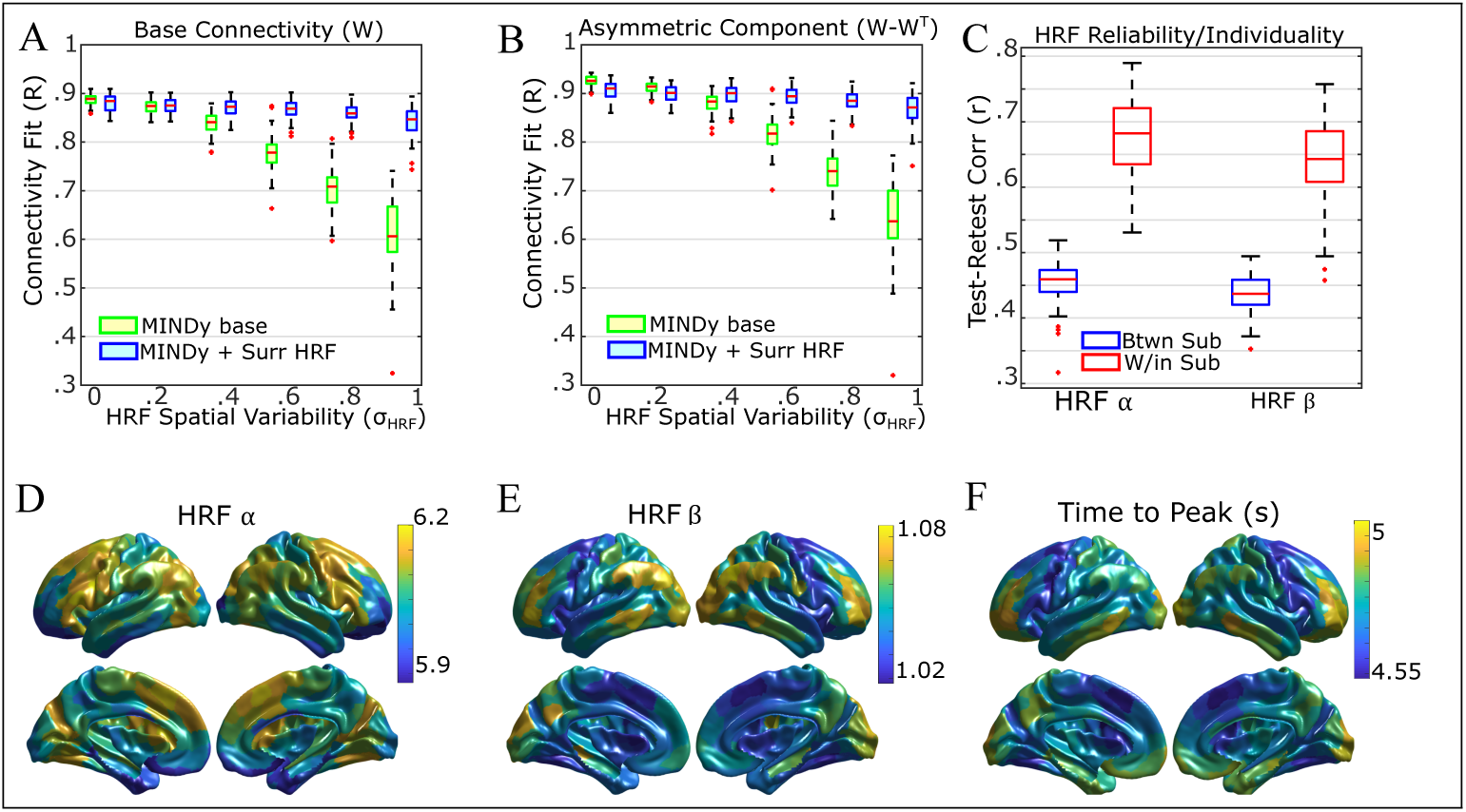
Incorporating HRF surrogate-deconvolution into MINDy. A) Without HRF modeling, connectivity estimates degrade with spatial variability in the neurovascular coupling. Fitting the HRF through surrogate deconvolution preserves performance. B) Same as (A) but for the asymmetric component of connectivity. C) HRF parameter estimates from HCP data are reliable across scanning days and subject-specific. D) Spatial map of the mean *α* parameter estimate across subjects. E) Same as (D), but for the second HRF parameter (*β*). F) Spatial map of the mean time-to-peak in the fitted HRF’s.

### E. Empirical Dual Estimation with the Human Connectome

Lastly, we tested the effects of using Surrogate Deconvolution in fitting MINDy models to data from the Human Connectome Project ([27]). By using empirical data, this analysis demonstrates that human hemodynamics are spatially variable and that accounting for this variability produces more nuanced and reliable brain models. Data consists of one hour of resting-state fMRI per subject spread across two days (30 minutes each). Data were processed according to the recommendations of Siegel and colleagues ([29]) and divided into 419 brain regions ([28]). We then fit MINDy models either with or without surrogate deconvolution to this data using the same fitting procedure and hyperparameters ([30]) as before. Results indicated the the parameters which describe the hemodynamic response function are reliable across scanning days and reliably differ between individuals (Fig. 4C). Each of the two HRF parameters had a stereotypical spatial distribution (Fig. 4D,E) as did the time-to-peak of the recovered HRF kernels (generated by substituting in the recovered kernel parameters). Time to peak was slowest for anterior prefrontal cortex, particularly in the right hemisphere (Fig. 4F). Because current knowledge of the “true” hemodynamic response is limited, future study establishing groundtruths for HRF variation across human cortex is needed to facilitate more rigorous empirical validation.

## V. CONCLUSION

Data-driven modeling remains one of the key challenges to neuroengineering and computational neuroscience. Although a wealth of theoretical model forms have been produced, the state-variables of these models (e.g. neuronal firing rate) are often difficult to directly measure *in – vivo* which complicates system-identification (model-parameterization). Instead, many clinical and experimental contexts record proxy variables which reflect the physiologically downstream effects of neuronal activity (e.g. on blood oxygenation, signaling molecules, and synaptic currents). In the current work, we aimed to parameterize conventional neural models using indirect measurements of neural activity. This problem involved simultaneously estimating the generative neural model as well as the latent neural activity thus comprising a dual-estimation problem. Through surrogate models, we approximated the state-estimation step as a parameterized deconvolution,, thus reducing computationally challenging dual-estimation problems to a closed-form, conventional identification problem. The primary advantage of this approach is speed/scalability.

Current approaches to model-based dual-estimation emphasize the joint/dual Kalman Filters and related Bayesian approaches (e.g. [16]). These approaches suffer, however, in terms of scalability and data quantity. As demonstrated in numerical simulations, the computational complexity of Kalman-Filtering limits application to relatively small models (Fig. 3F), whereas Surrogate Deconvolution enables optimization techniques that scale well with the number of parameters (Fig. 3F). However, despite requiring orders of magnitude fewer computations, Surrogate Deconvolution performs competitively with Kalman Filtering in estimating system parameters (Fig. 3B,C) as well as estimating states (latent neural activity; Fig. 3D,E). Thus, the computational advantages of Surrogate Deconvolution do not compromise accuracy.

Scalability is particularly salient in empirical neuroimaging, as several recent approaches have eschewed detailed modeling of physiological measurements (e.g. [30], [31]) in order to increase the spatial coverage of models. However, ground-truth simulations indicate that these reductions potentially compromise accuracy (Fig. 4 A,B). By contrast, methods augmented with Surrogate Deconvolution maintained high levels of performance (accuracy) even for extreme spatial variation in physiological signals. Interestingly, this variation appeared to be a reliable feature in empirical data with consistent differences across individuals (Fig. 4 C) and brain regions (Fig. 4 D-F) which can potentially lead to systematic biases (as opposed to random error) when these features are not modeled. Thus, for neuroimaging in particular, it may be critical to parameterize both the generative neural model and the measurement models to account for these biases. Surrogate Deconvolution provides a means to parameterize such models without compromising the detail of either component.

## Acknowledgments

MS was funded by NSF-DGE-1143954 from the US National Science Foundation. TB acknowledges R37 MH066078 from the US National Institute of Health. SC holds a Career Award at the Scientific Interface from the Burroughs-Wellcome Fund. Portions of this work were supported by AFOSR 15RT0189, NSF ECCS 1509342 and NSF CMMI 1537015, NSF NCS-FO 1835209 and NIMH Administrative Supplement MH066078-15S1 from the US Air Force Office of Scientific Research, US National Science Foundation, and US National Institute of Mental Health, respectively.

## VI. APPENDIX

### A. “Local” Field Potential Simulations

The “local” field potential recordings from Sec. IV-A were simulated using the discrete-time neural mass model:

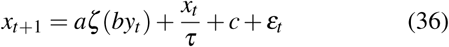

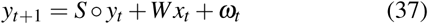

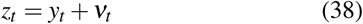

The paramaters *a* = 3, *b* = 1*/*5, and *τ* = 2 were fixed. For each simulation, the remaining neural-mass model parameters were redrawn from fixed, independent distributions: 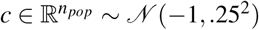 and *S* ∼ *𝒩* (.5, .2^2^) *∩* [.2, .8]. The connectivity parameter *W* was sampled using a two-step procedure:

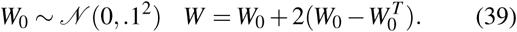

This exaggerated asymmetry serves to ensure solutions have nontrivial dynamics in the absence of noise. The noise processes *ε*_*t*_, *ω*_*t*_, and *v*_*t*_ were all independent, white Gaussian processes with the same variance for each population. For each simulation the standard deviations of *ε*_*t*_, *ω*_*t*_, and *v*_*t*_ were drawn from .25 + .5| *𝒩* (0, 1)|, .05 + .1|*𝒩* (0, 1)|, and .1 + .2|*𝒩* (0, 1)|, respectively. The variances assumed by Kalman Filtering were .5, .1, and .2 for *ε*_*t*_, *ω*_*t*_, and *v*_*t*_, respectively.

### B. Randomized Networks and MINDy Hyperparameters for simulated fMRI

Ground-truth simulations for BOLD fMRI (Sec. IV-D) were produced by a 40 node Hopfield-type ([25]) recurrent neural network with asymmetric connectivity:

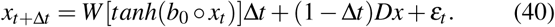

Here, the timescale of integration was Δ*t*=.1s and measurement occurred every 700*ms*. The process noise *ε*_*t*_ was Gaussian (*σ* ^2^ = .625) and independent between channels. The simulation parameters and generic MINDy fitting hyper-parameters were generally identical to those in the original 40-network MINDy simulations ([30]). Ground-truth connectivity parameters (*W*) for the simulations were generated by a hyperdistribution characterized by four hyperparameters which scale the reduced-rank magnitude (*σ*_1_), sparseness (*σ*_2_), degree of asymmetry (*σ*_*a*_), and degree of population clustering 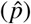. These hyperparameters are distributed *σ*_1_, *σ*_*a*_ ∼ *𝒩* (4,.1^2^) and *σ*_2_ ∼ *𝒩* (3,.1^2^). The hyperparameter 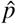 is either 1 or 2 with equal probability. These parameters were used to generate three matrices (*M*_1_, *M*_2_, *M*_3_) distributed as follows:

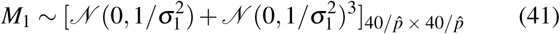

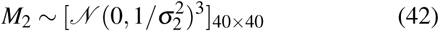

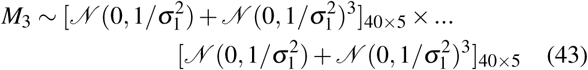

To generate population clustering we use the ones matrix 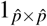 and define 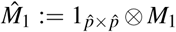 in which ⊗ denotes the Kronecker product. The final connectivity matrix (*W*) for each simulation is formed as follows:

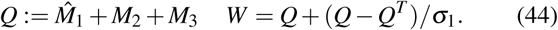

The slope vector *b*_0_ ∈ R^40^ is distributed *b*_0_ ∼ *𝒩* (6, (.5)^2^) and the diagonal decay matrix *D* has (diagonal) elements iid. distributed *D*_*i,i*_ ∼ *𝒩* (.4, .1^2^) *∩* [.2, ∞]. Deconvolved time series were z-scored. Base MINDy regularization parameters for the 40-node simulation were generated by rescaling the empirical fMRI regularization parameters 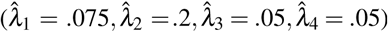 by 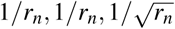 *and* 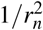 respectively with *r*_*n*_ = 10 is the approximate ratio between the number of empirical brain regions (419) and those used in the simulation (40) ([30]). The maximum-rank of the low rank component *W*_*L*_ was 15. Initial values for MINDy parameters were distributed as in ([30]). The NADAM update rates for the HRF parameters *α* and *β* were 5 × 10^−4^ and 2.5 × 10^−4^, respectively for the 40-node simulation. Surrogate deconvolution used the third-order bivariate polynomial basis {1, *α, β, α*^2^, *αβ, β* ^2^, *α*^3^, *α*^2^*β, αβ* ^2^, *β* ^3^} which was fit to the z-scored deconvolution surfaces.

### C. Surrogate-HRF MINDy Parameters for HCP Data

For the empirical data, MINDy used the original regularization parameters 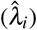. NADAM update rates were 2.5 × 10^−4^ for *α* and 2.5 × 10^−5^ for *β*. Data was preprocessed according to Siegel and colleagues ([29]) and smoothed via nearest-neighbor. Deconvolution was performed using Wiener’s method with noise-signal-ratio =.002. On each minibatch, next-step predictions were made for 250 sequential frames using an HRF kernel length of 30 TRs (21.6s) and parameter updates were performed using NADAM for 6000 minibatches. As before, surrogate deconvolution used the third-order bivariate polynomial basis {1, *α, β, α*^2^, *αβ, β* ^2^, *α*^3^, *α*^2^*β, αβ* ^2^, *β* ^3^} to approximate the z-scored deconvolved time-series. For fitting surrogate coefficients, *α* was assumed uniform on [5, 7] and *β* was assumed uniform on [.5, 1.5]. Expected values were taken by sampling this two-dimensional space along an evenly-spaced 10 × 10 rectangular grid.

